# 5-HT4 receptor agonists treatment reduces tau pathology and behavioral deficit in the PS19 mouse model of tauopathy

**DOI:** 10.1101/2023.02.03.526871

**Authors:** Shan Jiang, Eric J. Sydney, Avery M. Runyan, Rossana Serpe, Helen Y. Figueroa, Mu Yang, Natura Myeku

## Abstract

Accumulation of tau in synapses in Alzheimer’s disease (AD) has been shown to cause synaptic damage, synaptic loss, and the spread of pathology through synaptically connected neurons. Synaptic loss correlates with a decline in cognition, providing an opportunity to investigate strategies to target synaptic tau to rescue or prevent cognitive decline. One of the promising synaptic targets is the 5-HT4 receptor present post-synaptically in the brain areas involved in the memory processes. 5-HT4R activation exerts synaptogenic and pro-cognitive effects involving synapse-to-nucleus signaling essential for synaptic plasticity. However, it is not known whether 5-HT4R activation has a therapeutic effect on tauopathy. The goal of this study was to investigate the impact of stimulation of 5-HT4R in tauopathy mice. Our results show that 5-HT4R agonism led to reduced tauopathy and synaptic tau and correlated with increased proteasome activity and improved cognitive functioning in PS19 mice. Thus, stimulation of 5-HT4R offers a promising therapy to rescue synapses from toxic synaptic tau.

## Introduction

Besides main features such as the deposition of amyloid beta (Aβ) plaques and neurofibrillary tangles (NFT), other common characteristics of AD are synaptic dysfunction, structural damages on the synapses, and synaptic loss (Peng et al., 2022). The synaptic changes occur in the early stages of the disease before the symptoms of memory dysfunction (Chen et al., 2018), with recent studies suggesting that synaptic tau accumulation promotes synaptic impairment that contributes to the cognitive deficit (Xia et al., 2017). Tau is predominantly an axonal protein, and its presence in the synaptic compartments is low in physiological conditions. However, early in AD pathogenesis, tau is mislocalized to synapses with high levels of hyperphosphorylated and high molecular weight (HMW) tau in the post-synaptic compartments (DeVos et al., 2018; Schaler et al., 2021). Consequently, tau pathology spreads through synaptically connected neurons following the progression of the disease as defined by Braak stages I-VI (Braak and Braak, 1991) and demonstrated by PET imaging studies in patients with AD (La Joie et al., 2020) and progressive supranuclear palsy (PSP) (Cope et al., 2018). Notably, symptoms of memory dysfunction and later cognitive impairment correlate with the propagation of tau pathology from the hippocampus to the cortex (Qian et al., 2017).

Interestingly, the accumulation of synaptic hyperphosphorylated and HMW tau correlates with the accumulation of synaptic ubiquitinated proteins in AD brains and mouse models (Schaler et al., 2021; Tai et al., 2012), suggesting a disruption of synaptic proteasome-mediated proteolysis, which can further lead to synaptic damage and synaptic loss. Indeed, recent studies have reported that inhibition of synaptic protein degradation machinery can lead to the accumulation of tau in dendrites and the loss of dendritic spines (Balaji et al., 2018). Furthermore, mechanistic studies have shown that oligomers and aggregates can reduce proteasome activity by impairing the function of ATPase subunits of the 26S proteasome, presumably by obstructing the gate opening of the 26S proteasome (Myeku et al., 2016) or acting as allosteric inhibitors and stabilizing the closed gate conformation of the proteasome (Thibaudeau et al., 2018).

The recent understanding that tau propagation across neurons involves synaptic secretion to the extracellular space and subsequent internalization by the neighboring neurons has led to the development of immunotherapy clinical trials against extracellular tau species. However, the recent failure of anti-tau antibody therapies may hint that clearance of extracellular tau pool may not be an effective strategy as most pathological tau is intracellular (Congdon et al., 2022). Thus, increasing the turnover of intracellular tau, especially enhancing proteasome activity, which is recognized as the cell’s first defense mechanism against accumulating proteotoxicity, has been suggested as a new or additional approach to delay the onset or lessen the pathology in proteinopathy disorders (Myeku and Duff, 2018). Enhancing proteasome activity could have therapeutic potential but is still a relatively unexplored field. To this end, our team (Myeku et al., 2016) has recently shown that hyperphosphorylation of the 26S proteasome by PKA can mediate enhanced proteasome activity, reduced tauopathy, and cognitive decline in a mouse model. Subsequently, our group has shown that enhancing degradation of toxic tau restricted to synapses via stimulation of PAC1R, a Gs-coupled GPCR present on the membrane of post-synaptic compartment reduced the spread of tau, extracellular tau seeds, and overall tau aggregation in mice (Schaler et al., 2021). GPCRs mediate critical physiological functions and are considered the most successful drug targets (34% of all FDA-approved drugs) prescribed for a broad spectrum of diseases (Hauser et al., 2017). The class of Gs-coupled GPCRs that mediate synapse-to-nucleus signaling by stimulating the AC/cAMP/PKA/CREB pathway has many therapeutic potentials specifically for synaptogenesis and synaptic plasticity (Fisher et al., 2016; Teich et al., 2015).

The GPCRs of the serotonergic innervation are considered the largest group of receptors found in mammals, with wide expression in the CNS and periphery (Smith et al., 2017). Of the 14 serotonin receptor subtypes identified in mammals, 5-HT4R is strongly linked to AD pathology, synaptic plasticity, and neuroprotection (Hannon and Hoyer, 2008). 5-HT4R is mainly an excitatory GPCR localized post-synaptically and is expressed in the brain areas involved in cognition and emotion, such as the hippocampus, prefrontal cortex, amygdala, and basal ganglia (Beliveau et al., 2017). The AD-like effects at the behavioral, cellular, and molecular levels have been reported in a 5-HT4R KO mouse (Karayol et al., 2021). Moreover, the depletion of the serotonergic system and 5-HT4R is evident in post-mortem AD brains (Reynolds et al., 1995).

A large body of animal studies has shown that 5-HT4R agonists appear to be central to enhanced memory function by synaptic plasticity-related enhancement. The activation of 5-HT4R stimulates adenylate cyclase (AC) to produce cAMP. The signal is amplified by the activation of PKA, which is known to enhance the cAMP response element-binding protein (CREB)-mediated transcription. Furthermore, activation of 5-HT4R leads to the release of the non-amyloidogenic N-terminal ectodomain of APP, known as sAPPα, which has a potent memory-enhancing effect by displaying neuroprotective and neurotrophic properties (Hashimoto et al., 2012). Importantly, 5-HT4R has been a successful drug target for the intestine to promote intestinal peristalsis (Lanthier et al., 2020). However, early FDA-approved agonists have had adverse cardiovascular events. The latest FDA-approved 5-HT4R agonist, prucalopride, is highly selective with an excellent safety profile and no adverse cardiovascular effects (Yiannakou et al., 2015).

In this study, we tested whether two highly selective 5-HT4R partial agonists, prucalopride and RS-67333, can enhance tau clearance in synapses via proteasome and attenuate tauopathy in PS19 mice. Furthermore, our transcriptomics analyses from AD postmortem brains show that transcripts related to the 5-HT4R signaling cascade were reduced in neurons but not in other cells across Braak stages and that the levels of 5-HT4R inversely correlated with Aβ plaques.

## Results and Discussion

### Chronic 5-HT4R stimulation reduced tau pathology

To investigate the effect of 5-HT4R stimulation on tauopathy, we used two selective, high-affinity 5-HT4R partial agonists with good brain penetration (Johnson et al., 2012), prucalopride (3 mg/kg) and RS-67333 (2 mg/kg). Prucalopride is an FDA-approved medication for treating constipation with an excellent safety profile and has recently been shown to exert a pro-cognitive effect and increase hippocampal function in a small clinical trial in healthy subjects (de Cates et al., 2021). In animal models, RS-67333 was extensively shown to have pro-cognitive (Quiedeville et al., 2015) and anxiolytic/anti-depressant-like activity (Mendez-David et al., 2014).

Adult 5-6 months old PS19 or wild-type (WT) mice were treated intraperitoneally twice daily for six weeks. At this age, PS19 mice develop early-stage tauopathy. After behavioral studies, cortical and hippocampal tissue was collected to generate the total and insoluble extracts to assess the hyperphosphorylated and aggregated tau species and to analyze the results by quantitative immunoblotting. Our data show that prucalopride had a significant effect in reducing phosphorylated and total tau forms in the total (**Fig. 1A**) and the insoluble brain extracts (**Fig. 1B**), likewise RS-67333 showed a similar effect (**Fig. 1A,B)**, albeit a non-significant trend was detected in the total extract for pS396/pS404, and total tau forms (**Fig. 1A**). Immunohistochemical analyses of the hippocampus, shown here in the CA1 region, also showed that 5-HT4R agonists markedly reduced pS202/pT205 tau epitope (**Fig. 1C**).

**Fig. 1.**
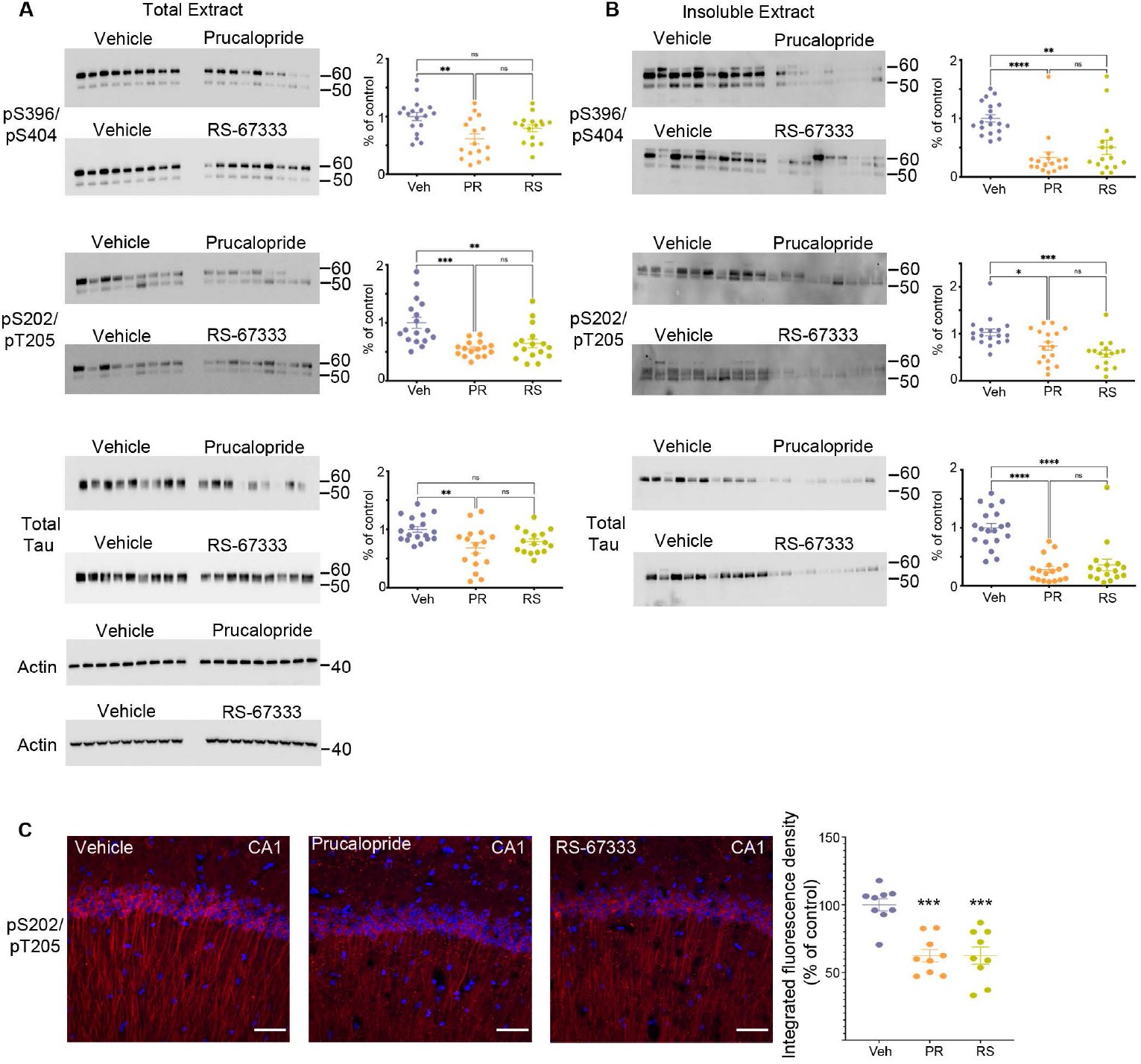
Stimulation of 5HT4R by prucalopride or RS-67333 attenuates tau pathology in PS19 mice. (**A** and **B**) Representative immunoblots and corresponding densitometric quantification for pS396/pS404, pS202/pT205 tau epitopes and total tau from (**A**) total extracts and (**B**) insoluble extracts from brains of PS19 mice treated with vehicle (Veh), prucalopride (PR) or RS-67333 (RS). Quantified densitometry for immunoblots expressed as percent relative to vehicle-treated mice. (**C**) Immunofluorescence labeling of pS202/pT205 tau epitope and quantification of integrated fluorescence density for pS202/pT205 tau epitope in the CA1 region of the hippocampus of PS19 mice treated with vehicle, prucalopride or RS-67333. Scale bar (100μm). Scatter plots represent the quantification of immunoreactivity normalized to actin. Statistical analyses of vehicle (n=19), prucalopride (n=17), and RS-67333 (n= 16) mice were performed in two sets (**Fig. 1** and **Supplementary Fig. 1 and 2**). Quantification of immunofluorescence intensity used three brain slices from 9 animals/groups. For statistical analyses, we used one-way ANOVA followed by Bonferroni multiple comparison post hoc tests. Error bars mean ± SEM. ns; not significant, *P < 0.05, **P < 0.01, ***P < 0.001, and ****P < 0.0001.

While the molecular signature of 5-HT4R stimulation in the CNS has been reported through the anti-amyloidogenic cascade (Cochet et al., 2013) and the synapse to nucleus pro-cognitive and neurotrophic effect (Hagena and Manahan-Vaughan, 2017), our data expand the therapeutic potential of the 5-HT4R by showing that 5-HT4R agonists treatment could also reduce tau burden and overall tauopathy in the brain.

### 5-HT4R stimulation reduced synaptic tau

Synaptic tau accumulation and trans-synaptic propagation of pathological tau contribute to synaptic degeneration and cognitive deficits in AD and other tauopathies. We have shown previously that post-synaptic accumulation of tau in the early stages of the disease is toxic and seed competent, able to template and cause aggregation of naïve endogenous tau before the neurofibrillary tangles appear in the brain (Schaler et al., 2021). Moreover, combating the accumulation of tau in the post-synaptic compartment by enhancing the turnover of tau restricted to synapses by the proteasome leads to reduced tauopathy (Schaler et al., 2021). In this study, we tested whether stimulation of 5-HT4R found predominantly on the surface of post-synaptic compartments can enhance proteasome activity to clear synaptic tau. We isolated gradient-purified synapses and subsequently separated isolated synapses into pre and post-synaptic fractions from the vehicle, prucalopride, and RS-6733 treated PS19 mice to detect tau changes within synapses by quantitative immunoblotting. Prucalopride treatment significantly decreased the total and pS396/pS404 tau in the pre and post synaptic compartments (**Fig. 2A,B,C**)

**Fig. 2.**
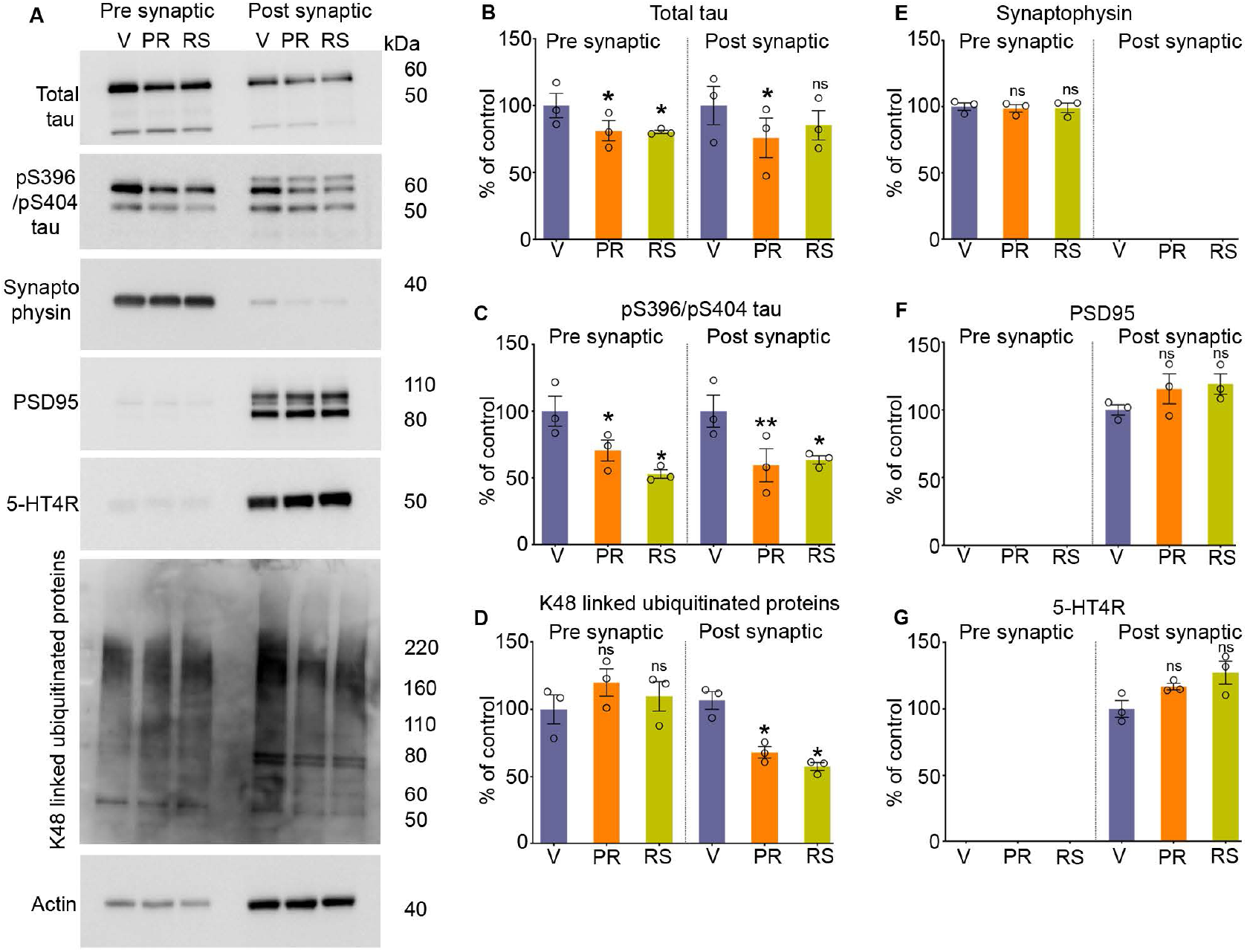
Stimulation of 5HT4R by prucalopride or RS-67333 reduces synaptic tau and ubiquitinated proteins in PS19 mice. (**A**) Representative immunoblots of pre-synaptic and post-synaptic fractions from the brains of PS19 mice treated with vehicle (V), prucalopride (PR), or RS-67333 (RS). Blots were probed for total tau, pS396/pS404, and pS202/pT205 tau epitopes, K48-linked ubiquitinated proteins, for pre-synaptic (synaptophysin) and post-synaptic (PSD95) markers, and 5-HT4R. Actin was a loading control and was used for normalization. Quantified densitometry for immunoblots, expressed as percent relative to vehicle-treated mice. Three fractionation experiments were performed, and for each fractionation experiment, four mouse hemi-cortices were pooled together (overall, n = 12 hemicortices per treatment). One-way ANOVA followed by Bonferroni, multiple comparisons post hoc tests, was conducted. Data are presented as means ± SEM; ns; not significant, *P < 0.05, **P < 0.01.

Similarly, RS-67333 showed a significant decrease in the pS396/pS404 tau in both synaptic compartments, and total tau was significantly decreased in the pre-synaptic fraction and showed a non-significant trend decrease in the post-synaptic compartment (**Fig. 2A,B,C**). While tau species were reduced, we did not detect any changes in the markers of the pre and post-synaptic compartments, synaptophysin (**Fig. 2A,E**) and PSD95 (**Fig. 2A,F**), respectively. In fact, PSD95 levels showed a moderate increase in the treatment group, with prucalopride or RS-67333, compared to vehicle control. We confirmed the presence of 5-HT4R in the post-synaptic compartment as it was reported to be present post-synaptically, and its levels were slightly increased in the agonist treatment, but the increase was non-significant (**Fig. 2 A,G**).

Next, we tested the status of K-48-specific ubiquitinated proteins exclusively turned over by the 26S proteasome. Our data show that the K-48 specific ubiquitinated proteins were significantly reduced in the post-synaptic fractions in prucalopride and RS-67333 treatment compared to the vehicle treatment group (**Fig. 2A,D**). The pre-synaptic fractions showed no changes in the ubiquitinated protein amount (**Fig. 2A,D**). Suggesting that proteasome activity was mainly enhanced in a spatially restricted manner in the post-synaptic compartments.

The findings here suggest that agonism of 5-HT4R, which is present predominately post-synaptically, can enhance the signaling cascade to mediate enhanced proteasome activity and reduce tau restricted to local stimulation. Over time enhanced tau turnover can lead to reduced tauopathy throughout the brain.

### 5-HT4R agonism enhanced proteasome activity

5-HT4R activation stimulates adenylate cyclase as a primary mode of signal transduction and increases cAMP concentration. The signaling cascade is amplified when cAMP interacts and activates PKA, which is known to enhance gene expression related to synaptic plasticity and neurotrophic factors by phosphorylating transcription factors, such as CREB. Moreover, a new function of PKA-mediated phosphorylation leads to hyperphosphorylation of the 26S proteasomes, conferring enhanced degradation capacity to proteasome complexes to degrade proteins in the cells, especially aggregating-prone proteins such as tau (Myeku et al., 2016), poly-GA (Khosravi et al., 2020), huntingtin (VerPlank et al., 2020), and TDP43 (Lokireddy et al., 2015). Altogether this raises the interest in targeting 5-HT4R, not only for its neurogenic and plasticity-related memory improvement but concomitantly as a mode of enhancing the turnover of misfolded toxic proteins that can initially accumulate within synapses. To this end, we assessed proteasome capacity to degrade fluorogenic substrate kinetically over 120 min (**Fig. 3A**). The slope of the reaction was calculated and showed that 5-HT4R agonists significantly increased the proteasome’s proteolytic capacity over time (**Fig. 3B**). The results suggest that 5-HT4R agonism is a positive regulator of proteasome activity via its canonical cAMP/PKA signaling, leading to reduced synaptic tau species and ubiquitinated proteins.

**Fig. 3.**
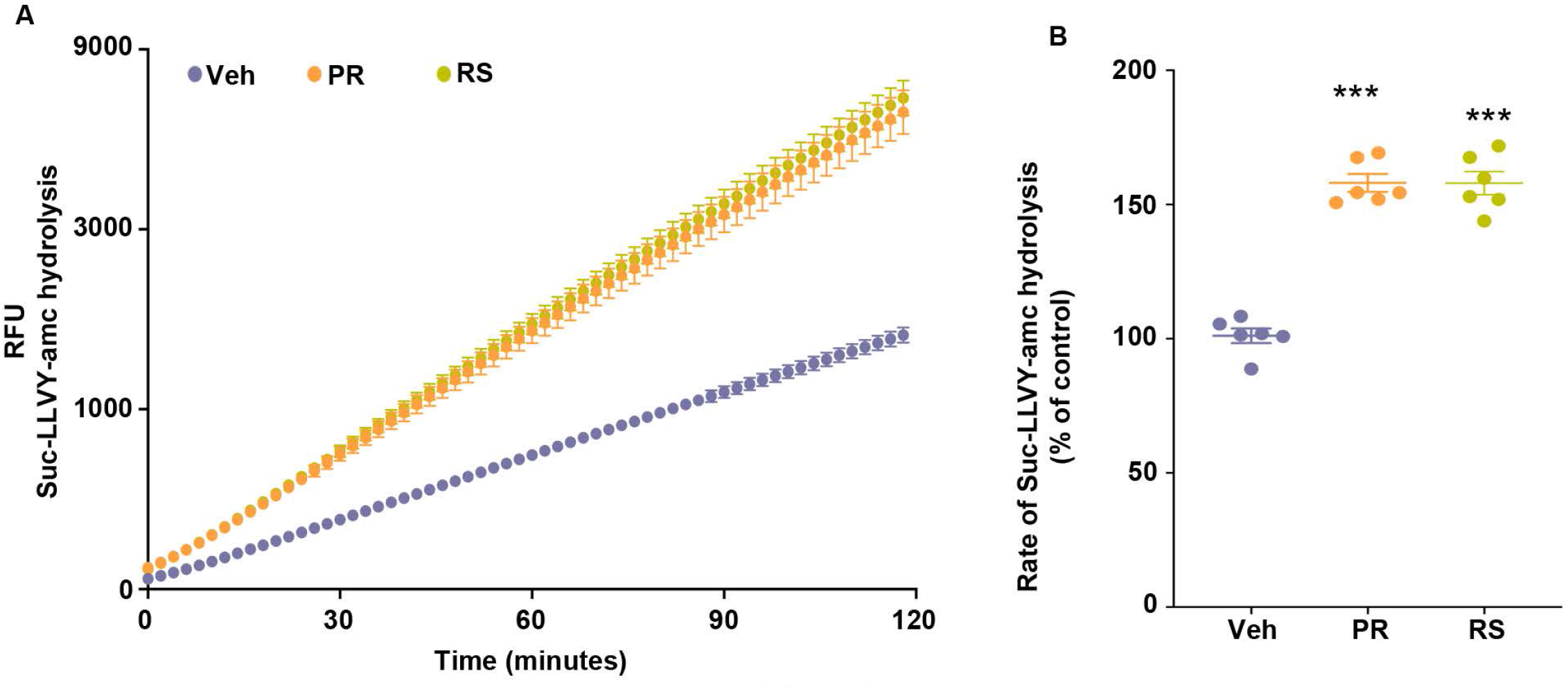
Stimulation of 5HT4R by prucalopride or RS-67333 enhances proteasome activity in the brain of PS19 mice. (**A**) The peptidase activity of proteasomes from the total brain extracts of PS19 mice treated with vehicle (V), prucalopride (PR), or RS-67333 (RS) was assessed for 120 min using the fluorogenic substrate Suc-LLVY-amc (40 μM). Y-axis represents relative fluorescence units (RFU). (**B**) The graph represents the rate of substrate hydrolysis by proteasomes from (A) expressed as a percentage of vehicle control. Total proteasome lysate from n=6 animals/treatment was used in the assays, and three independent proteasome kinetic assays were performed. One-way ANOVA, followed by Bonferroni multiple comparison post hoc tests, was used for (B). Error bars mean ± SEM. ***P < 0.001.

### 5-HT4R agonism reduced anxiety-related behavior and improved cognitive performance in early-stage tauopathy

PS19 mice exhibit mild behavioral phenotypes. Therefore, many standard behavioral tests do not show any changes in cognitive performance in these mice. This is true for other tauopathy models with mild pathology. Recently a detailed characterization of PS19 mice across ages has shown that hippocampal-dependent impairment in spatial reference memory was only apparent in female mice at age 12 months (Sun et al., 2020). However, at this age, these mice develop hindlimb paralysis due to transgene expression in the spinal cord and die between 10 and 12 months of age (Jankowsky and Zheng, 2017). For our studies, we performed the open field test, which evaluates spontaneous locomotor activities as well as anxiety-like behaviors (Kraeuter et al., 2019), and a non-maternal nest-building test, sensitive to hippocampal damage to evaluate cognition (Sun et al., 2020). P301S mice from three treatment groups (vehicle, prucalopride, and RS-67333, 6-7 months old) showed a similar decrease in the ambulatory distance over 60 minutes (**Fig. 4A**). However, when we separated the treatment groups by sex, vehicle-treated females showed significantly increased total ambulatory distance (**Fig. 4B,D),** indicating enhanced hyperactivity, compared to male vehicle-treated PS19 littermates. Treatment with agonists blunted the longer traveled distance in the female group (**Fig. 4B,D**), normalizing the results to the male PS19 littermates (**Fig. 4C,D**), which was similar in all treatment groups. In WT mice (6-7 months old), there was no difference in sex and treatment effect across the three groups (**Fig. 4 E-H**).

**Fig. 4.**
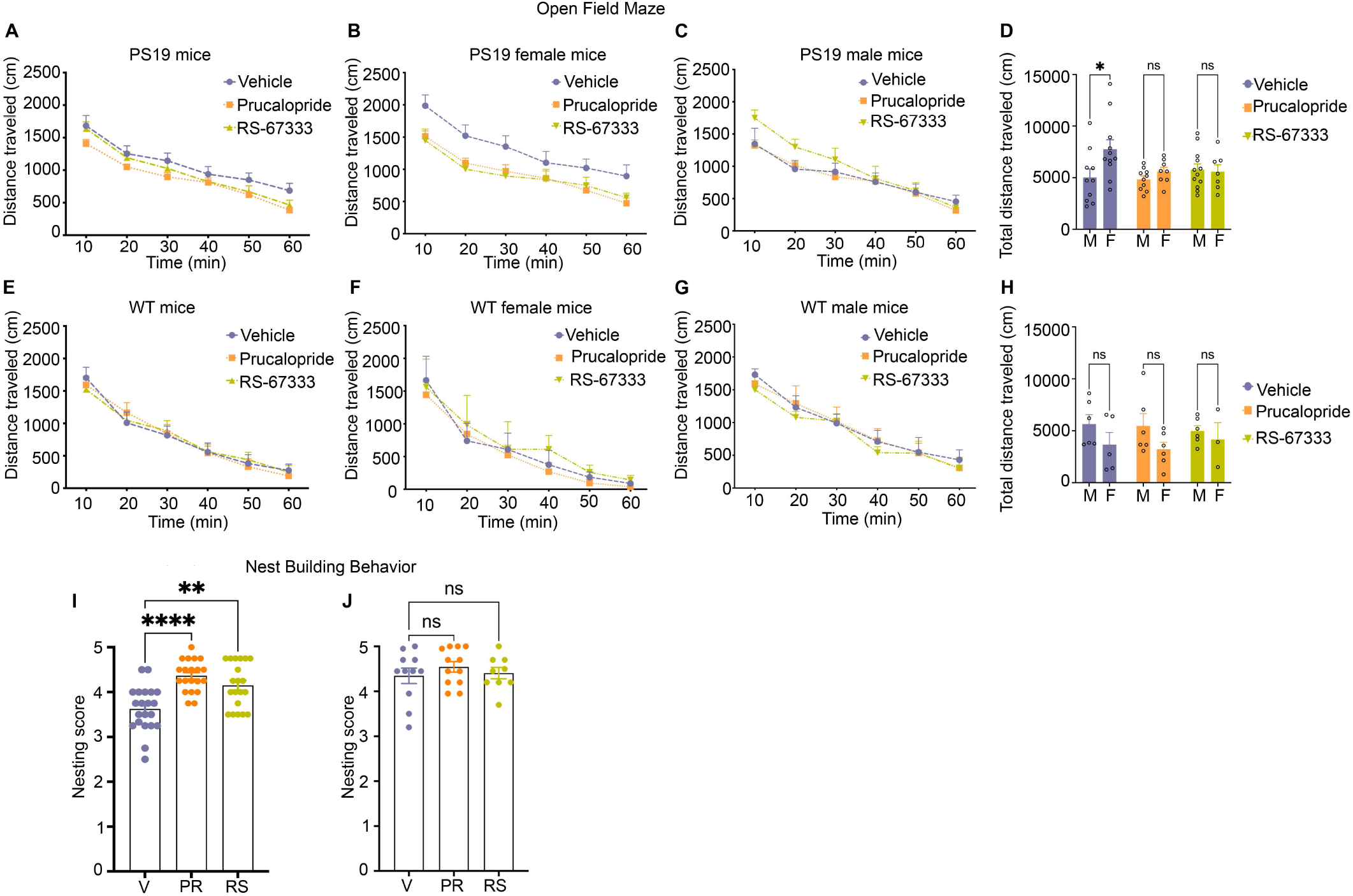
Stimulation of 5HT4R by prucalopride or RS-67333 reduces anxiety-related behavior and improves cognitive performance in early-stage tauopathy in PS19 mice. **(A-H**) The open field test was used to assess locomotor activities and anxiety-related behaviors by measuring the ambulatory distance over 60 min. PS19 and wild-type mice were treated with vehicle, prucalopride, or RS-67333. Vehicle (n = 21), prucalopride (n = 18), RS-67333 (n = 19) in the PS19 mouse group, and vehicle (n = 11), prucalopride (n = 12) and RS-67333-(n = 9) in the wild-type mouse group. (**A**) The traveled distance of PS19 mice showed no significant changes due to the treatment effect. (**B,C**) Grouped by sex and treatment. (**B**) Vehicle-treated females showed significantly longer total ambulatory distance compared to female littermates treated with prucalopride or RS-67333. (**C**) Male littermates showed no changes in ambulatory distance due to treatment. (**D**) Total ambulatory distance in a vehicle, prucalopride, or RS-67333 treated groups separated by sex, shows that vehicle-treated female PS19 mice exhibit significantly higher ambulatory distance compared to their vehicle-treated male littermates. Whereas prucalopride or RS-67333 treatment normalized the total distance of the female group to the distance traveled by male littermates. (**E-H**) wild-type (WT) mice showed no difference in ambulatory distance in sex and treatment effect across the three groups. The open-field test data were analyzed using a two-way repeated-measures ANOVA with post hoc Bonferroni correction. (**I,J**) Nest building test for PS19 and WT mice. (**I**) Nesting scores from vehicle (n = 21), prucalopride (n = 18), RS-67333 (n = 19) in PS19 mice. (**J**) Nesting scores from vehicle (n = 11), prucalopride (n = 12), RS-67333 (n = 9) in WT mice. Nest building test scores were assigned 24 hours after one compressed cotton nestlet was placed in each cage. Nesting data were analyzed using the non-parametric Kruskal–Wallis test. Data are reported as means ± SEM. n.s., not significant; **P < 0.01 and ***P < 0.001.

Furthermore, the non-maternal nest-building performance showed that vehicle-treated PS19 mice exhibited significantly impaired nest-building performance compared to prucalopride and RS-67333 treated groups (**Fig. 4I**). In contrast, the WT mice showed no treatment effect in nest-building performance across the groups (**Fig. 4J**). Although vehicle-treated female PS19 mice exhibited hyperactive locomotor activities compared to male vehicle-treated littermates, they did not perform worse in the nest-building test (data not shown). Overall, the behavioral testing shows that 5-HT4R agonists normalized the anxiety like-behavior in female PS19 mice, which could also be related to the anxiolytic effect of 5-HT4R agonism. And that hippocampal-related behavioral task was improved with both 5-HT4R agonists.

### 5-HT4R/AC/PKA/CREB genes are decreased with exacerbated tau aggregation in human post-mortem brains

5-HT4R plays an essential role in processes related to learning and memory mediated by the dominant synapse to nucleus signaling cascade through AC/cAMP/PKA pathway, modulating the expression of plasticity/learning-related proteins such as BDNF, AKT, and CREB (Roux et al., 2021). To assess the mRNA expression levels of the canonical proteins in this cascade (5-HT4, Gsα, Gβ, Gγ, AC, PKA R, and C subunits and CREB), we performed bioinformatics analyses from bulk and single nucleus RNA sequencing from available datasets. Bulk tissue sequencings were available from Mount Sinai Brain Bank (MSBB) study with 299 samples, the Religious Orders Study/Memory and Aging Project (ROSMAP) study with 633 samples, and the Mayo study with 319 samples. Single nucleus RNA sequencing (sn-RNAseq) were available from Mathys *et al*. (Mathys et al., 2019) which comprised 80,660 single-nucleus transcriptomes, and Seattle Alzheimer’s Disease Brain Cell Atlas (SEA-AD), which comprised 1,240,908 single-nucleus transcriptomes.

Before performing the analyses, we assessed the tissue expression specificities of these genes and chose the gene subtypes with high specificity in the brain (Fagerberg et al., 2014).

RNA-seq of bulk tissues presented by heat map shows that except *CREB1*, all genes in the pathway were decreased predominately in advanced Braak stages consistently in all the three bulk tissue cohorts (MSBB, ROSMAP, and Mayo) (**Fig. 5A-C**). Because multiple brain regions were sequenced for the same individuals in the MSBB cohorts, the parahippocampal gyrus (PHG), which has an early onset of AD pathology, was chosen to be presented (**Fig. 5A**). Similarly, the other brain regions, such as a frontal pole, inferior frontal gyrus, and superior temporal gyrus, showed the same expression patterns of these genes as in PHG (data not shown). Meta-analysis results by combining the three cohorts are shown in **Supplementary Fig. S3**. Except for *CREB1*, the other genes in the pathway show decreased expression in advanced Braak stages. Furthermore, we examined the relationship between the *HTR4* gene and Aβ plaque load from the MSBB cohort where Aβ plaque load information is available and found a statistically significant negative correlation between the HTR4 expression and the Aβ plaque load (**Fig. 5D**). While consistent, bulk RNA seq results could reflect tissue degeneration in late stages of the disease. Thus, for our next step analyses, we used snRNA-seq datasets from Mathys *et al*.,(Mathys et al., 2019) and from the Seattle Alzheimer’s Disease Brain Cell Atlas (SEA-AD) with the aforementioned single nucleus counts, respectively. Due to the undetectable low expression levels of some genes in sn-RNAseq, which are known as dropouts, and the relatively low expression of the genes of interest, the gene expression in the single cell heatmaps was represented and visualized as expression percent instead of the mean value (Muus et al., 2021; Qiu, 2020). Our analyses show that the expression percent of all the HTR4 pathway-related genes, including *CREB1*, were significantly reduced with advanced Braak stages in excitatory and inhibitory neurons (**Fig. 5E,F**).

**Fig. 5.**
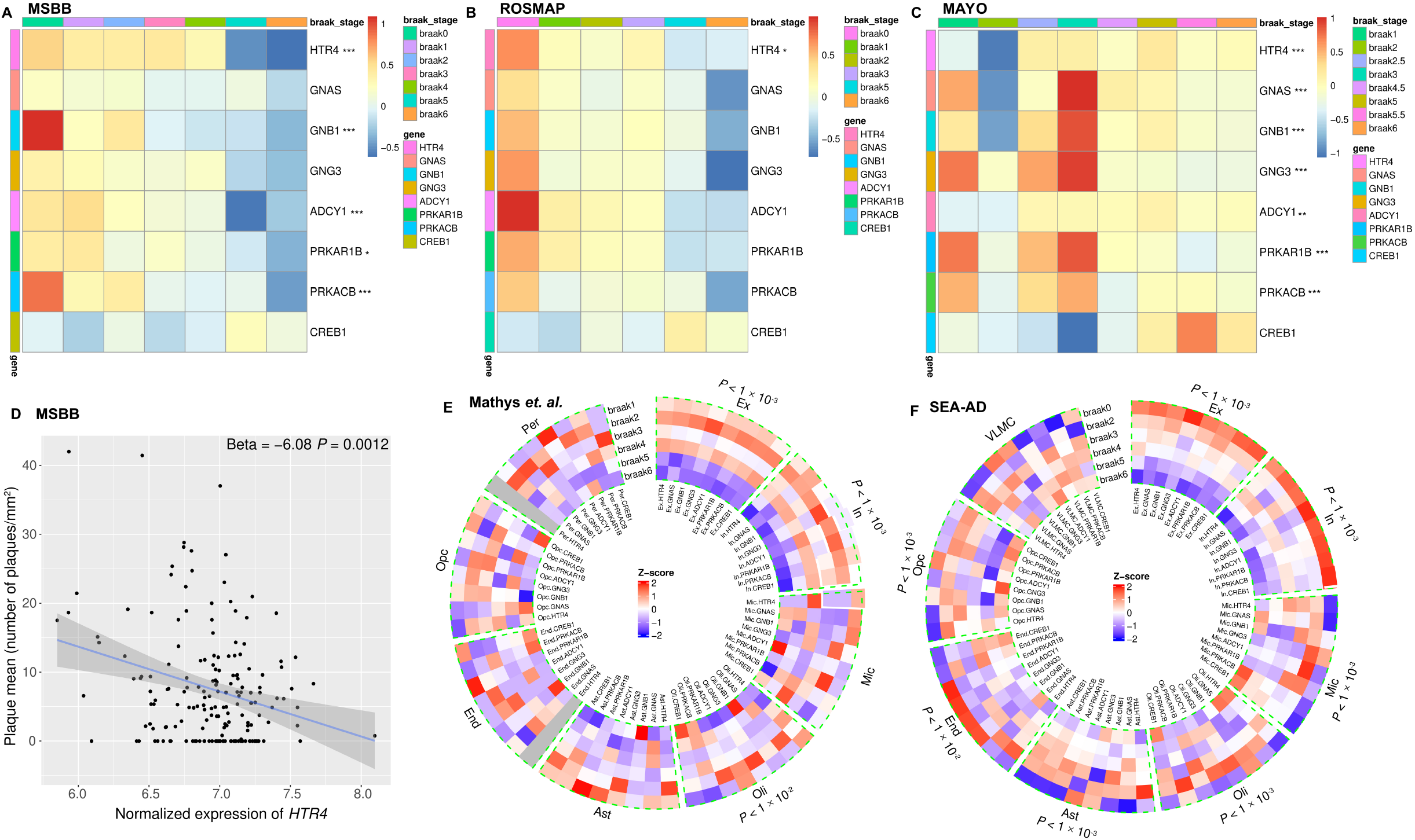
The RNA seq analyses show decreased expression of 5-HT4R/AC/PKA/CREB1 pathway-related genes with exacerbated tau aggregation. (**A-C**) Bulk RNA-seq data analyses presented as heatmaps showing HTR4/AC/PKA/CREB1 pathway-related genes (except CREB1) were decreased in advanced Braak stages in the AD post-mortem brains. Brain areas (**A**) parahippocampal gyrus (PHG) from MSBB study. (**B**) Dorsolateral prefrontal cortex from the ROSMAP study. (**C**) Temporal cortex from the Mayo study. Gene expressions were scaled with Z-score transformation. Ordinal regression was used to assess the correlation between Braak stages and gene expression after adjusting for the effects of confounding factors such as age, gender, ethnicity, PMI, RIN and library size. *P < 0.05, **P < 0.01, and ***P < 0.001 after FDR correction. (**D**) Scatter plot showing the HTR4 expression was negatively correlated with Aβ plaque measure in PHG from MSBB study with linear regression after adjusting for confounding effects. (**E,F**) Single nucleus RNA-seq analyses presented in circular heatmaps show the expression percent of HTR4/AC/PKA/CREB1 pathway-related genes were decreased with advancing Braak stages in excitatory and inhibitory neurons. (**E**) Prefrontal cortex from Mathys et al. dataset study. (**F**) The middle temporal gyrus from SEA-AD studies and Gene expression percentages were scaled with Z-score transformation. Sn-RNA-seq data were analyzed using the Kruskal-Wallis test; *P < 0.05, **P < 0.01, and ***P < 0.001 after FDR correction. Ex, (excitatory neurons); In, (inhibitory neurons); Mic, (microglia); Oli, (oligodendrocytes); Ast, (astrocytes); End, (endothelial cells); Opc, (oligodendrocyte progenitor cells); Per, (pericytes); VLMC, (vascular leptomeningeal cells).

For the non-neuronal cell types, in the Mathys et. al. study, the genes showed no specific expression patterns with developing tau pathology (**Fig. 5E**), which may be attributable to low cell counts in the dataset. However, in SEA-AD, which sequenced over one million cells, the selected genes in microglia and astrocytes were significantly increased with advanced Braak stages (**Fig. 5F**).

By comparing the expression patterns of *CREB1* in bulk RNA-seq and snRNA-seq, it was clear that the increased *CREB1* expression in bulk RNA-seq (**Fig. 5A-C**) in advanced Braak stages were due to the increased expression in microglia, oligodendrocytes, endothelial cells, and oligodendrocyte progenitor which counteracted the decreased expression in neurons (**Fig. 5E,F**). The bioinformatics results show that in AD, the 5-HT4R serotonergic pathway is downregulated predominantly in neurons, reflecting the occurrence of serotonergic degeneration in AD brain pathology (Smith et al., 2017). Thus, preservation of 5-HT4R and its signaling cascade by agonists starting from the asymptotic stages of the disease could delay cognitive decline and tau aggregation in AD.

## Conclusion

Synaptic dysfunction and loss due to the accumulation of tau in synapses and subsequent trans-synaptic propagation of tau pathology underlines the importance of designing synaptic therapy approaches to preserve synapses and halt the spread of tauopathy and memory decline.

Our study tested one such approach using the receptor-mediated enhanced tau clearance in synapses in early tauopathy in PS19 mice. We had previously shown that activating a peptidergic Gs-GPCR, the PAC1R present in synapses could stimulate cAMP/PKA-mediated proteasome activity and enhance the turnover of toxic synaptic tau and overall tauopathy (Schaler et al., 2021). However, peptide-related therapies have limited clinical application due to rapid enzymatic degradation and diminished concentration in the brain. To overcome these limitations, in the present study, we targeted the 5-HT4 serotonergic receptor, a successful druggable target that is a promising target for the prevention of MCI in AD patients (de Cates et al., 2021). 5-HT4R activation with small molecule agonists leads to synaptogenic, neurotrophic, and neuroprotective effects through well-documented mechanisms of production of non-amyloidogenic sAPPα, activation of cAMP/PKA/CREB signaling, and enhanced BDNF expression. However, while the 5-HT4R stimulation outcomes on the above pathways have been well documented, there are no data reported on the effect of 5-HT4R activation on tau pathology. Here we tested the long-term administration of the 5-HT4R partial agonists, prucalopride, and RS 67333 on the impact on the tauopathy mouse model. We show that activation of 5-HT4R leads to reduced synaptic tau and overall tauopathy. This is correlated with enhanced proteasome degradation capacity and reduced ubiquitinated proteins in synapses. Moreover, the behavioral testing showed that 5-HT4R agonism had an anxiolytic effect on female PS19 mice and improved hippocampal-related behavioral tasks, providing further support for a pro-cognitive effect of 5-HT4R agonism. Altogether, this raises the interest in targeting 5-HT4R in preserving synapse-to-nucleus serotonergic pathways, which are downregulated in AD, maintaining synapses integrity by reducing the toxic accumulation of synaptic tau and enhancing plasticity-related memory as a crucial path to identifying viable targets for AD treatment.

## Materials and Methods

### Mice

The heterozygous PS19 (P301S) transgenic line and the WT mice with the same background (B6C3) were used with equal male and female mice. The transgene is driven by the murine prion protein (Prnp) promoter and expresses the P301S human 1N4R tau isoform. Mice were housed in pathogen-free conditions under a normal 12-h light/dark cycle.

### Intraperitoneal injections

PS19 and WT mice aged 5-6 months were randomly assigned to three experimental groups; vehicle, prucalopride (3mg/kg), and RS-67333 (2mg/kg); and treated twice daily intraperitoneally for six weeks with the vehicle and 5-HT4R highly selective partial agonists; prucalopride (Sigma, cat# 1371) or RS 67333 (Sigma, cat# 1882). The drugs were dissolved in vehicle (0.9% NaCl and 1% DMSO). Drug administration and testing were performed in three sets of mice (two sets of PS19 mice and one set of WT mice). Animal experiments were in full compliance with the US National Institutes of Health Institutional Animal Care and Use Committee guidelines and overseen by Columbia University Irving Medical Center.

### Immunofluorescence

Mouse brains were isolated after transcardial perfusion with PBS, and half of the brains were drop-fixed in 4% PFA overnight and then subjected to cryoprotection treatment in 30% sucrose in PBS for 24 h. The other hemisphere was used for homogenization assays. Free-floating brain sections (50 μm) from brains sectioned in the sagittal plane were used. The sections were incubated at 4 °C overnight with primary antibody diluted in PBS containing 0.3% Triton X-100 and 5% normal goat serum blocking solution (Vector Laboratories, #S-1000). Anti-mouse monoclonal pS202/pT205 1:1000 (AT8, 1:500, #MN, 1020 from Thermo Fisher) was used. Following washes, sections were incubated with goat anti-mouse IgG Alexa 594 (Thermo Fisher #A-11005, 1:500). Nine animals (3 slices each) per treatment were used. Staining was visualized by confocal microscopy, Zeiss LSM710 confocal microscope at 20× dry objective. Sequential tile scans were performed to capture images of 1,024 × 1,024 resolution. All images from the same experiment were taken at the same laser intensity and detector gain.

### Kinetic assay for proteasome activity

Cortical brain tissue was harvested and homogenized in a buffer containing 50 mM Tris-HCl (pH 7.4), 5 mM MgCl2, 5 mM ATP, 1 mM dithiothreitol, 1 mM EDTA, 10 mM NAF (Sodium fluoride), 25 mM β-glycerolphosphate, phosphatase inhibitors, and 10% glycerol, which preserved 26S proteasome assembly and centrifuged at 20,000 x g for 25 min at 4°C. The supernatant was normalized for protein concentration determined by Bradford assay. The kinetic assay was used to measure proteasome activity. Samples (lysate, 25 μg per well) and proteasome substrate, 100 μM Suc-LLVY-amc were mixed, reading fluorescence signal Ex. 380 nm; Em. 460 nm over 2 hours.

### Tissue fractionation and protein extraction

Frozen hemispheres free of cerebellum and brainstem were weighed and homogenized without thawing in RIPA buffer (10× volume/weight) (50 mM Tris-HCl, pH 7.4, 1% NP-40, 0.25% sodium deoxycholate, 150 mM NaCl, 1 mM EDTA, 1 mM phenylmethylsulfonyl (PMSF), 1 mM sodium orthovanadate, 1 mM sodium fluoride (NaF), 1 μl/ml protease inhibitor mix. Homogenates were centrifuged for 10 min at 3,000 × g at 4 °C. Protein assay was performed on the clear supernatants representing the total extract used to analyze the total protein levels. Sample volumes were adjusted with RIPA buffer containing 100 mM DTT and NuPAGE LDS Sample Buffer 4× buffer (Life Technologies) and boiled for 5 min. The sarkosyl-insoluble extracts, which are highly enriched in aggregated tau species, were generated when 250 μg aliquots from the total protein extracts were normalized into 200 μl final volume containing 1% sarkosyl, followed by ultra-centrifugation at 100,000 × g for 1 h at 4°C. Without disturbing the pellet, the supernatant was transferred to new tubes. The pellet was resuspended in 100 μL RIPA buffer containing DTT and NuPAGE LDS Sample Buffer 4× buffer, followed by vortexing for 1 min and 5 min heating at 95 °C.

### Immunoblot analysis

Samples (2.5–10 μg protein) were typically run on 4–12% Bis-Tris gels (Life Technologies; WG1403BOX10) using MOPS buffer (NP0001). Proteins were analyzed after electrophoresis on SDS-PAGE and transferred onto 0.2-μm nitrocellulose membranes. Samples were run independently for each protein. i.e., blots were not stripped and re-probed with different antibodies as most proteins had similar molecular weights. Blots were blocked and incubated with primary and secondary antibodies at the below concentrations. Membranes were developed with enhanced chemiluminescent reagent (Immobilon Western HRP substrate and Luminol reagent (WBKLS0500, Millipore) using a Fujifilm LAS3000 imaging system. ImageJ (http://rsb.info.nih.gov/ij) was used to quantify the signal. Relative intensity (fold change or fold increase, no units) is the ratio of the value for each protein to the value of the respective loading control.

### Antibodies

Monoclonal human tau (CP27; 1:5000) and pS396/pS404 (PHF1; 1:2500) were generous gifts from the late Dr. Peter Davies. Mouse monoclonal pS202/pT205 Tau (AT8, 1:2500, #MN, 1020) was from Thermo Fisher. Monoclonal anti-rabbit K-48 linked ubiquitin (1:1000, D905) from Cell Signaling. Mouse monoclonal anti-PSD-95 (1:5000, 6G6-1C9, ab2723); rabbit monoclonal synaptophysin (YE269, ab32127, 1:5000) were from Abcam. Loading control anti-actin (1:5000 AC-74, #A2228) was from Sigma. Secondary antibodies were from Jackson Immunoresearch, anti-mouse (115-035-003), and anti-rabbit (111-035-003)

### Synaptic fractions

#### Fractionation assay

Four hemi cortices/experiment from mice (2 hemi cortices/gender) were gently homogenized in ice-cold homogenization buffer (320 mM sucrose made in hypotonic buffer, 25mM HEPES-KOH pH 7.5, 1mM EDTA, 5mM MES pH 7.5, 5mM MgCl2, 5mM ATP pH 7.5, 1mM DTT, 10mM NAF, 25mM βglycerol phosphate, phosphatase, and protease inhibitors). The homogenates were centrifuged at 500 x g for 5 min at 4°C. The supernatant (total extract) was further centrifuged at 19, 000 x g for 20 min at 4°C. The resulting supernatant is a cytosolic fraction, and the pellet is a crude synaptosome resuspended in homogenizing buffer.

#### Discontinuous sucrose density gradient

Crude synaptosome extracts were layered on top of a nonlinear sucrose gradient (1.2M, 0.8M, and 0.32M sucrose from bottom to top) and centrifuged at 300,000 x g for 4 hours at 4°C. After which, synaptosomes sediment at the interface between 1.2M and 0.8M sucrose layer. To separate synaptosomes into pre and post -synaptic fractions, collected synaptosomes (~3–4 mL/sample) were diluted in 0.01 mM CaCl2, 20mM Tris–HCl pH 6, 1% Triton X-100, with protease and phosphatase inhibitors and mixed by inversion for 20 min at 4°C. After incubation, samples were centrifuged at 40,000 × g for 20 min. The resulting pellet was collected as a post-synaptic fraction, and the supernatant was collected as a pre-synaptic fraction. After separating pre and post-synaptic compartments, fractions were precipitated in ice-cold acetone and concentrated by Amicon-15 10kDa cut-off tubes. Both fractions were resuspended in PBS with 0.05% Triton X-100 with protease/phosphates inhibitors and sonicated. The fractionation assay was repeated at least three times. Protein concentration was determined by Bradford assay. Samples were diluted to 0.5 μg/ul and 2.5 μg/well loaded for western blotting. Immunoblots for each protein were run independently at an equal amount of proteins. For quantification, the relative values of three independent fractionation experiments from the vehicle groups were normalized to 100 percent and were compared to the treatment group. The values were then plugged into GraphPad Prism 9 software to generate bar graphs with S.E.M.

### Nest building test

The nest-building behavior has been used to assess mice cognition. PS19 or WT mice were singly housed and were given intact compressed cotton nestlets in the center of the cage. After 24 hours, images were taken for each nest from corresponding mice. Nest scoring used a 1-5 scale, as described previously (Neely et al., 2019). Scores: 1-untouched nestlet; 2 - partially torn nestlet, 3-partially complete nest, 4-nest was almost formed, 5 nest complete. All mice were tested after 6 weeks of treatments.

### Open field test

The Open Field is the most commonly used test for spontaneous exploratory activity in a novel environment, incorporating measurements of locomotion and anxiety-like behaviors. The Open Field test was performed following previously described protocols (Cho et al., 2021). Exploration was monitored during a 30 min session with Activity Monitor Version 7 tracking software (Med Associates Inc.). Briefly, each mouse was gently placed in the center of a clear Plexiglas arena (27.31 x 27.31 x 20.32 cm, Med Associates ENV-510) lit with dim light (~5 lux), and is allowed to ambulate freely. Infrared (IR) beams embedded along the X, Y, Z axes of the arena automatically track distance moved, horizontal movement, vertical movement, stereotypies, and time spent in center zone. Data are analyzed in six, 10-min time bins. Arenas are cleaned with 70% ethanol and thoroughly dried between trials

### Human transcriptomic analyses of HTR4/AC/PKA/CREB1 pathway-related genes

All the human datasets for bulk RNA transcriptomic data on AD were accessed from the AD Knowledge Portal (https://adknowledgeportal.synapse.org/). We examined the three cohorts, Mount Sinai Brain Bank (MSBB), Religious Orders Study/Memory and Aging Project (ROSMAP), and Mayo study. MSBB includes 299 individuals with multiple brain region tissue samples (214 parahippocampal gyrus samples, 261 frontal pole samples, 221 inferior frontal gyrus samples, and 240 superior temporal gyrus samples), and ROSMAP includes 633 individual dorsolateral prefrontal cortex samples. Mayo study comprises 355 individuals with 319 temporal cortex (TCX) and 278 cerebellum samples. Because the cerebellum is less affected in AD, especially in the early stage, the 278 cerebellum samples from Mayo were removed from the examination.

Raw transcriptomic count data were normalized with variance stabilizing transformation with DESeq2 (Love et al., 2014) The effects of HTR4/AC/PKA/CREB1 pathway-related gene expression levels on the degree of tau aggregation, which was categorized by Braak scores, were examined with ordinal regression after adjusting for the confounding effects. For MSBB, the confounding effects from age, gender, ethnicity, post-mortem interval (PMI), RNA integrity number (RIN) and library size were adjusted. For ROSMAP, the confounding effects from age, gender, ethnicity, education, PMI, RIN and library size were adjusted. For Mayo TCX samples, because all the samples were from the white population, the confounding effects from age, gender, RIN and library size were adjusted. The false discovery rate (FDR) approach was applied to multiple testing corrections. Then meta-analyses were performed to integrate the regression results across the three cohorts on individual genes with R package meta (Balduzzi et al., 2019).

### Human single nuclei RNAseq analyses of HTR4R/AC/PKA-CREB1 pathway-related genes

Single nucleus RNA-seq (snRNA-seq) data from Mathys et al., which consists of 80,660 single-nucleus transcriptomes from the prefrontal cortex (PFC) of 48 individuals from ROSMAP, was accessed from the AD Knowledge Portal (Mathys et al., 2019) snRNA-seq data from Seattle Alzheimer’s Disease Brain Cell Atlas (SEA-AD), which is comprised of 1,240,908 single-nucleus transcriptomes from the middle temporal gyrus (MTG) of 84 individuals, was accessed from Chan Zuckerberg Initiative Science (https://cellxgene.cziscience.com/collections/).

Raw snRNA-seq count data were normalized with log2 (TP10K + 1), where TP10K,or transcripts per ten thousand, is the RNA-seq quantification measure that accounts for differences in sequencing depth across cells. A mixed-effect regression model with individual samples as a fixed effect is not applicable to examine the effect of the expression of genes on the degree of tau aggregation due to the dropouts in snRNA-seq (Qiu, 2020). To determine whether the expression of HTR4/AC/PKA/CREB1 pathway-related genes differs among different degrees of tau aggregation for each cell type, a non-parametric one-way analysis of variance (ANOVA) and Kruskal-Wallis test was applied followed by post hoc pairwise comparisons with Wilcoxon rank sum test. The false discovery rate (FDR) approach was applied to multiple testing corrections

### Statistical and bioinformatics analyses

For mouse data,statistical analyses for Figs. 1–4 were performed with Prism 9 (GraphPad Software, San Diego, CA). P < 0.05 was considered significant. Data were assessed for normality using the Shapiro-Wilk test used to determine the homogeneity of variance. Data for Figs 1 and 3 were analyzed using one-way ANOVA with Bonferroni post hoc correction. Data for Fig 2 were analyzed with one-way and two-way ANOVA with post hoc Bonferroni correction. Data for Fig 4 (OF test) were analyzed with two-way repeated measure ANOVA with post hoc Bonferroni correction. Data for nest building results (Fig.4) were analyzed with the Kruskal Wallis test with multiple comparisons of Dunn’s correction.

Statistical and bioinformatics analyses for human data for Fig. 5 and Supplementary Fig. 3 were performed with R (version 4.2.2) and python (version 3.9.13). For Fig. 5A–5C, the effects of HTR4/AC/PKA/CREB1 pathway-related gene expression levels on the degree of tau aggregation were examined with ordinal regression after adjusting for the confounding effects, such as age, gender, ethnicity, PMI, RIN and library size, with post hoc FDR correction. For Fig. 5D, the effect of HTR4 gene expression on Aβ plaque measure in PHG from MSBB study was examined with linear regression after adjusting for the confounding effects from age, gender, ethnicity, PMI, RIN and library size. For Fig. 5E–5F, due to the zero inflation introduced by undetectable dropouts in snRNA-seq, mixed-effect regression model was not applicable. Instead, we examined the differences of HTR4/AC/PKA/CREB1 pathway-related gene expression among different degrees of tau aggregation for each cell type with Kruskal-Wallis test followed by post hoc pairwise comparisons with Wilcoxon rank sum test with FDR correction. We used a relatively more liberal multiple test correction than the Bonferroni used in mouse data because of large sample sizes in bulk-seq data and large cell counts in snRNA-seq data. The heatmaps for Fig. 5A–5C were generated with pheatmap (version 1.0.12). The scatter plot for Fig. 5D was generated with ggplot2 (version 3.4.0). The circular heatmaps for Fig. 5E–5F were generated with circlize (version 0.4.15). The meta-analysis forest plots for Fig. S4 were generated with meta (version 6.0.0).

## Online supplemental material

Supplementary Figure 1

Supplementary Figure 2

Supplementary Figure 3

## Acknowledgments

This study was supported by grants from the National Institutes of Aging: AG070075, AG064244, AG055694 to N.M. Author contributions: N.M. designed the study. N.M. and S.J. wrote the manuscript. S.J. performed bioinformatics analyses. E.J.S., A.M.R. and R.S. performed experiments and generated the data. H.Y.F. assisted with and cared for mice. M.Y. is a director of the behavioral core at CUIMC and contributed to behavioral studies. All authors discussed the results and commented on the manuscript.

## Data availability

Bulk-seq data from MSBB (https://www.synapse.org/#!Synapse:syn20801188); Bulk-seq data from ROSMAP (https://www.synapse.org/#!Synapse:syn23650893);

Bulk-seq data from Mayo (https://www.synapse.org/#!Synapse:syn5550404); SnRNA-seq data from Mathys et al. (https://www.synapse.org/#!Synapse:syn18485175);

SnRNA-seq data from SEA-AD (https://cellxgene.cziscience.com/collections/1ca90a2d-2943-483d-b678-b809bf464c30).

