## supplemental material for "5-HT4 receptor agonists treatment reduces tau pathology and behavioral deficit in the PS19 mouse model of tauopathy"

**Fig. S1.** 5-HT4 receptor agonists treatment attenuates tauopathy- Study 1.
**Fig. S2.** 5-HT4 receptor agonists treatment attenuates tauopathy- Study 2.
**Fig. S3.** Meta-analysis by forest plots combining the three bulk RNA-seq datasets.

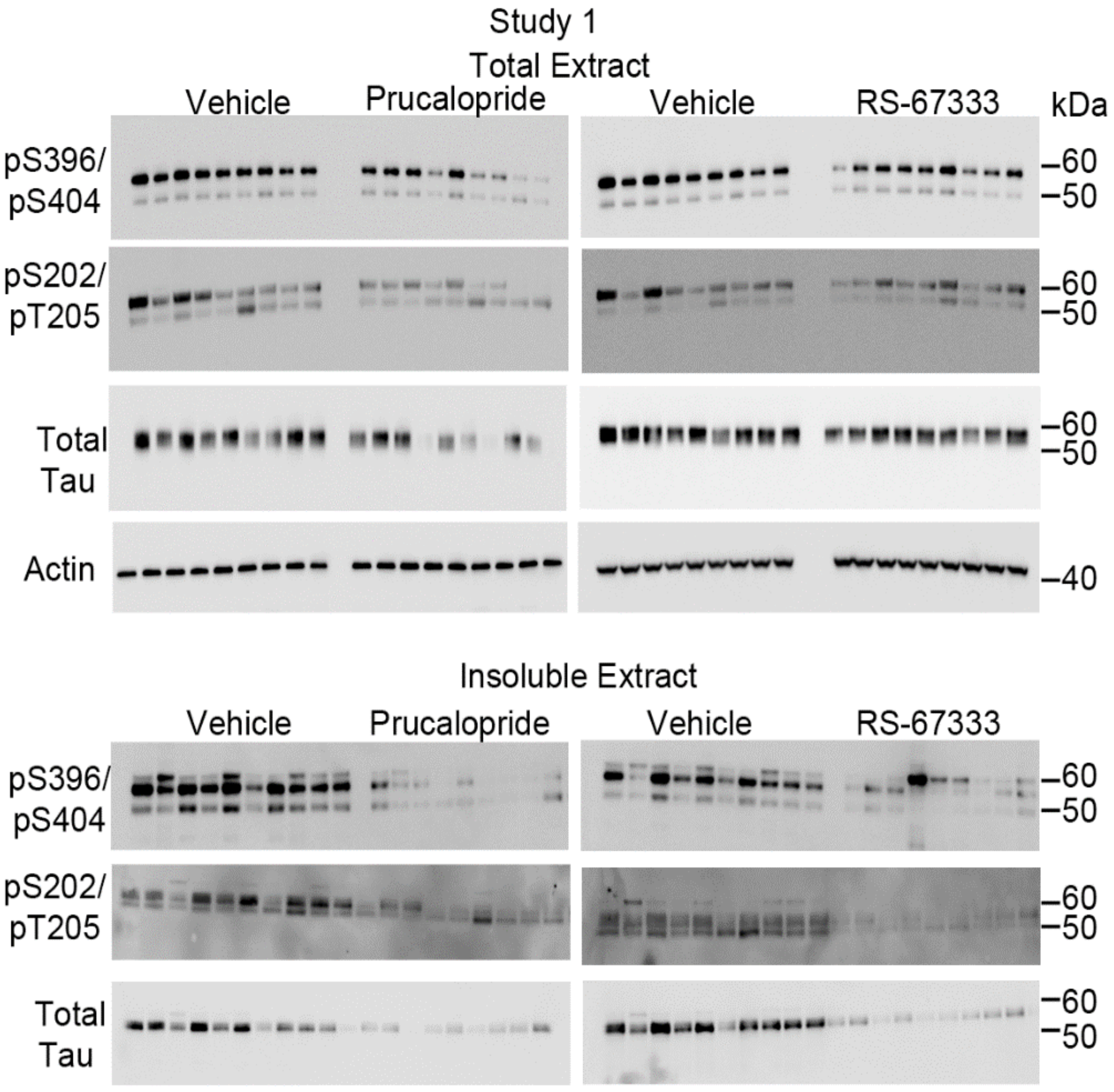


Supplementary Figure 1

**Fig. S1. 5-HT4 receptor agonists treatment attenuates tauopathy- Study 1.**
Summary of immunoblotting results from Fig.1. In vivo treatments were carried out in two sets/studies.

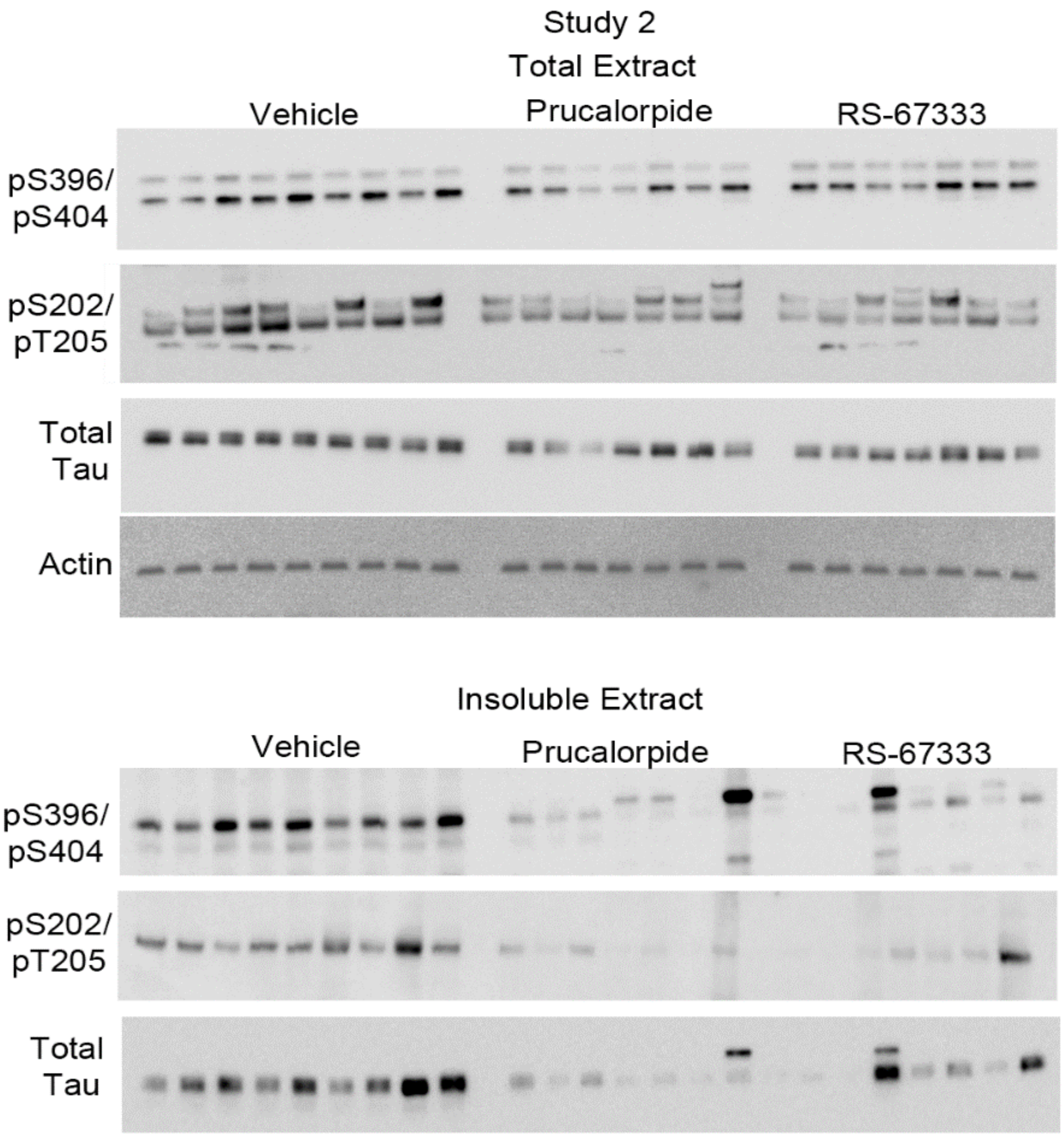


Supplementary Figure 2

**Fig. S2. 5-HT4 receptor agonists treatment attenuates tauopathy- Study 2.**
Summary of immunoblotting results from Fig.1. In vivo treatments were carried out in two sets/studies.

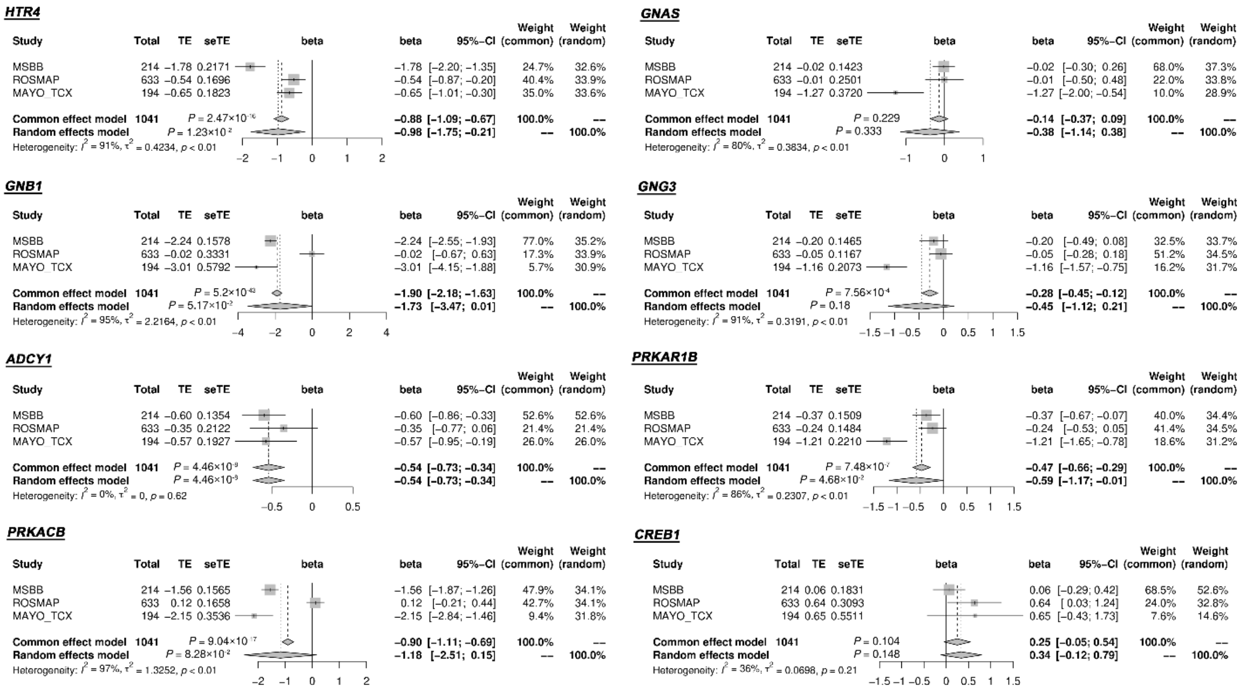


*Supplementray Figure 3*

**Fig. S3. Meta-analysis by forest plots combining the three bulk RNA-seq datasets.**
Data are point estimates of beta coefficients and a range of two-sided 95% confidence intervals (CI) of the beta coefficients. Heterogeneity among datasets was assessed with $I^2$ index. Combined beta coefficients and the corresponding P values were calculated in both fixed and random effect models.
